# Hsa_circ_0001947 suppresses acute myeloid leukemia progression by sponging hsa-miR-329-5p and regulating CREBRF expression

**DOI:** 10.1101/770420

**Authors:** Fengjiao Han, Chaoqin Zhong, Wei Li, Ruiqing Wang, Chen Zhang, Xinyu Yang, Chunyan Ji, Daoxin Ma

**Affiliations:** Department of Hematology, Qilu Hospital, Shandong University, Jinan 250012, China

**Keywords:** circRNAs, AML, hsa_circ_0001947, biomarker, hsa-miR-329-5p, CREBRF, therapy

## Abstract

Acute myeloid leukemia (AML) is a heterogeneous group of diseases resulting from clonal transformation of hematopoietic precursors through the acquisition of chromosomal rearrangements and multiple gene mutations. Accumulating evidence has indicated that aberrantly expressed circular RNAs (circRNAs) are involved in cancer development and progression. However, their clinical values and biological roles in AML remain unclear. In this study, we identified the aberrantly down-regulated profile of hsa_circ_0001947 in AML through microarray analysis and validated it with quantitative reverse transcription polymerase chain reaction (qRT-PCR). Then, we explored the clinical significance, biological functions and regulatory mechanisms of hsa_circ_0001947 in AML patients. The results showed that lower hsa_circ_0001947 expression was positively correlated with higher leukemia cells in bone marrow or peripheral blood, indicating poor prognosis. Further, bioinformatics analysis demonstrated hsa_circ_0001947-hsa-miR-329-5p-CREBRF network. Down-regulation of hsa_circ_0001947 by siRNA promoted cell proliferation, inhibited apoptosis, reduced drug resistance of AML cells, and also decreased the expression of its targeted gene, CREBRF. The mimics of hsa-miR-329-5p reduced drug resistance and decreased the expression of CREBRF, while its inhibitor manifested anti-leukemia effects and increased CREBRF expression. In vivo studies revealed that silencing hsa_circ_0001947 promoted the tumor growth in BALB/c nude mice. Collectively, our findings suggest that hsa_circ_0001947 functions as a tumor inhibitor to suppress AML cell proliferation through hsa-miR-329-5p/CREBRF axis, which would be a novel target for AML therapy.

## Introduction

Acute myeloid leukemia (AML) is characterized by blocked accumulation of clonal myeloid progenitor cells and exhibits complex clinical and biological heterogeneity[1]. Although great efforts have made in AML, the prognosis of AML is still poor and its 5-year survival rate is only 25%[2]. It is of clinical significance to explore the pathogenesis and identify new targets for AML. Circular RNAs (circRNAs) are a widely expressed class of non-coding RNAs generated in a diverse set of eukaryotic organisms, including animals, plants, yeasts and protists[3–5]. Recently it has been considered as the latest research hotspot in the field of RNAs[6]. Unlike the traditional linear RNAs (containing 5’and 3’ ends), circRNAs have a closed continuous loops, which are difficult to be degraded by RNA exonuclease and are more stable in expression [6–8]. The most common method of circRNAs biogenesis is through backsplicing, in which the 5’ splice site (5’ss) of a downstream exon is paired with the 3’ss of an upstream exon[3]. Backsplicing can occur in both linear pre-mRNAs and within lariat intermediates harboring skipped exons[9, 10]. CircRNAs were reported to be cell-specific, which are relatively abundant in the brain and exist in human body fluids such as blood[11]. The relationship between circRNAs and linear RNA isoforms was complicated. There are reports that circRNAs are produced in competition with linear RNAs[12]. However, it was also reported that there is no definite correlation between circRNAs and liner RNAs[13]. CircRNAs contain plenty of microRNA (miRNA) binding sites, which offer itself an important role as the miRNAs sponge in cells. Through the function of binding certain miRNAs, circRNA could eliminate the inhibitory effect of miRNAs on its target genes, thereby increasing the expression level of the target genes[7]. This mechanism is called competitive endogenous RNA (ceRNA)[7, 8]. Moreover, circRNAs are suggested to carry out diverse functions including sequestration and trafficking of proteins, regulation of transcription and generation of short proteins [14–16].

By interacting with disease-associated miRNAs, circRNAs play a significant regulatory role in various diseases, from which the tumor-related research caught our great attention. It has been reported that circTADA2As serve as miR-203a-3p sponge to suppress breast cancer progression and metastasis[17], and circPSMC3 is a sponge of miR-296-5p to suppress the proliferation and metastasis of gastric cancer by acting as a ceRNA[18]. In addition, circOMA1 mediated the miR-145-5p suppresses of tumor growth in nonfunctioning pituitary adenomas[19]. These findings indicate that circRNAs could be a promising new diagnostic markers and treatment targets in solid tumors.

However, there are rare data about the role of circRNAs in hematological diseases, especially in the AML until now. It was reported that silencing of circ_0009910 inhibits acute myeloid leukemia cell growth through increasing miR-20a-5p[20]. Moreover, we have proven that hsa_circ_0004277 could be characterized as a new biomarker and treatment target for AML via circRNAs profile and bioinformatics analysis. In particular, we found chemotherapy could significantly restore the expression of hsa_circ_0004277, indicating the increasing level of hsa_circ_0004277 was associated with successful treatment[21]. However, the detailed mechanism of circRNAs in the pathogenesis of AML is still unclear.

To better demonstrate the expression profile of circRNAs in AML, we performed a circRNA microarray in AML patients and healthy controls and presented an AML-specific circRNA feature. We identified that the hsa_circ_0001947 was one of the critical circRNAs that are frequently down-regulated in the bone marrow of AML patients. Then with the help of bioinformatics analysis, a detailed circRNA-miRNA-mRNA interaction network was described for hsa_circ_0001947. What’s more, in vitro and in vivo experiments showed that hsa_circ_0001947 could inhibit the progress of AML. Overall, our study demonstrates that hsa_circ_0001947 exerts critical regulatory role in the AML and may serve as a potential therapeutic target for AML.

## Materials and methods

### Patients and sample collection

BM samples from 106 AML patients and 15 healthy controls were obtained following informed consent at Qilu Hospital, Shandong University. The study was conducted in accordance with the Declaration of Helsinki, and the protocol was approved by the Ethics Committee of Qilu Hospital. Bone marrow mononuclear cells (BMMCs) from BM aspirates were separated by Lymphocyte Separation Medium (TBD, China) density gradient centrifugation. BMMCs from 6 AML patients and 4 controls were used for the circRNA microarray, and all the 120 samples were used for further detection or functional investigation.

### Cell culture

THP-1 and 293T cells were commercially obtained from the Cell Bank of Type Culture of the Chinese Academy of Sciences (Shanghai, China). Cells were cultured in RPMI-1640 (Gibco, USA) supplemented with 10% fetal bovine serum (FBS, Biological Industries, Beit HaEmek, Israel) in a humidified atmosphere at 37°C and 5% CO_2_.

### QRT-PCR

Total RNAs from 121 samples were extracted using Trizol (Ambion, Carlsbad, CA, USA) and were detected using a Nanodrop 1000 spectrophotometer (Thermo Scientifific, Waltham,MA, USA). And then total RNAs were reversely transcribed into cDNA using MLV RTase cDNA Synthesis Kit (Takara, Japan). Quantitative RT-PCR was conducted in Roche Applied Science LightCycler 480II Real-time PCR systems (Roche Applied Science, Indianapolis, IN, USA) in accordance to the manufacturer’s instructions. A comparative cycle threshold (Ct) method was used to analyze the gene expression level. GAPDH was used as an internal control. Primer sequences were presented in Table 1.

**Table 1.**
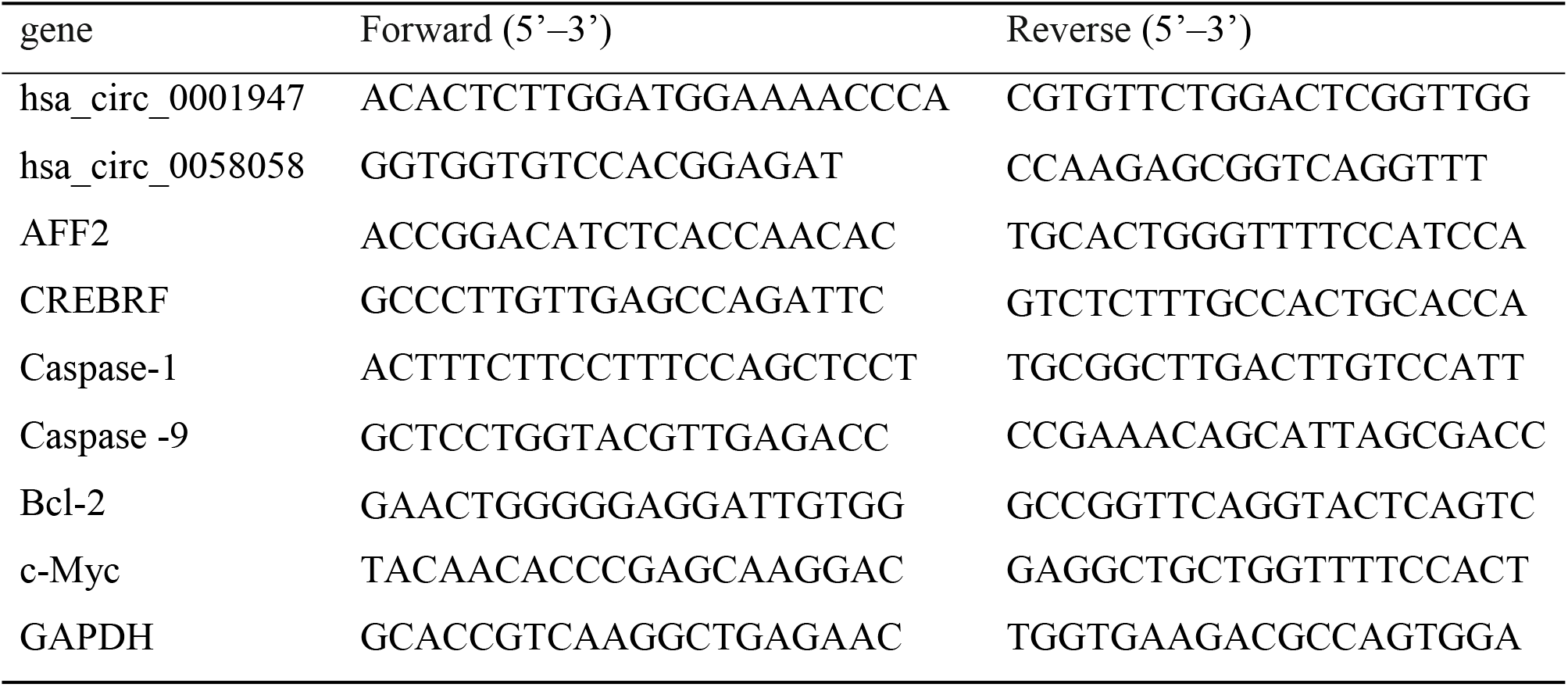
Primers used for qRT-PCR.

### RNase R treatment

Total RNA (1 μg) of THP-1 was mixed with 4 U RNase R at 37°C for 1 h. To assess the stability of hsa_circ_0001947 and line AFF2 mRNA, the expression levels were determined by using RT-PCR and agarose gel electrophoresis.

### Oligonucleotide transfection

The sicirc, hsa-miR-329-5p mimics, hsa-miR-329-5p inhibitor and their related control oligonucleotides were designed and synthesized by GenePharma (Shanghai, China). The sequences were presented in Table 2. All transfections were dissolved in deionized water to 20 uM according to the instructions. We transfected THP1 cells in 500ul medium with 2 ul transfections and 1 ul ExFect 2000 Transfection Reagent in 24-well plates.

**Table 2.**
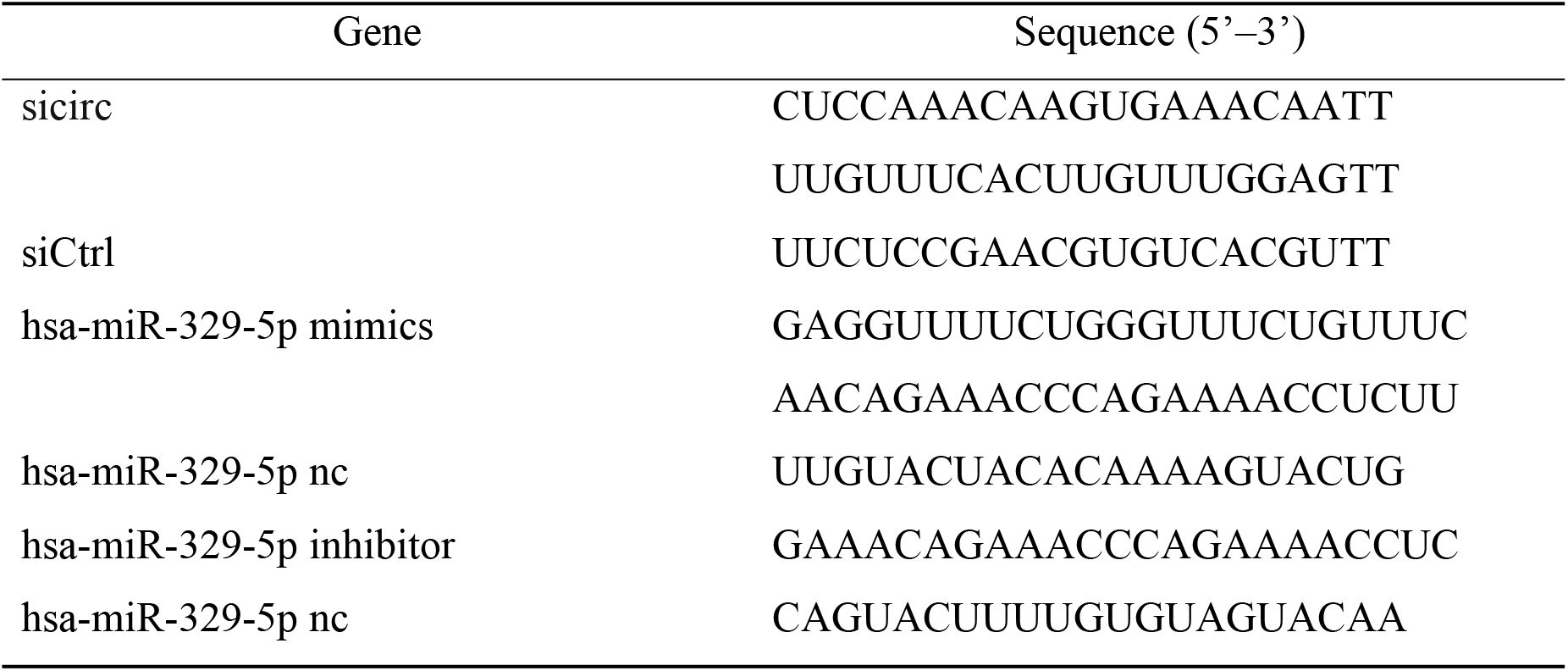
The sequences for oligonucleotide transfection.

### Cell proliferation assays

Cell proliferation was detected by cell counting kit-8 (CCK-8) assays according to the manufacturer’s protocols. THP-1 cell was planted into 96-well plates at 2500 cells/well. After different treatments, 10 ul of CCK-8 solution (BestBio, Shanghai, China) was added to each well and cells were incubated at 37°C atmosphere for 3 h, then the absorbance at 450 nm-630 nm-OD by microplate reader was measured following the instructions of the manufacturer (BioTek, USA).

### Apoptosis assay

Cell apoptosis was detected using a PI/Annexin V-FITC apoptosis detection kit (BestBio, Shanghai, China) according to the manufacturer’s instructions. Briefly, THP-1 cells transfected by sicirc, hsa-miR-329-5p mimics or inhibitor were treated with cytosine arabinoside (Ara-c) at 2 uM in 6-well plates for 48 h. They were harvested and re-suspended in 400 ul of Annexin V binding buffer and 5 μl of Annexin V-FITC in the dark at 4°C for 15 min. Next 10 μl of PI was added to the suspensions, and the cells were incubated in the dark at 4°C for 5 min. Then the samples were detected by Beckman Gallios cytometer. The final test results were subsequently analyzed with Kaluza software (Beckman Coulter, USA).

### Dual-luciferase reporter assay

A wild-type or mut-hsa-miR-329-5p fragment was constructed and inserted downstream of the luciferase reporter gene of the pmirGLO Vector (Promega, USA). We used ExFect 2000 Transfection Reagent to transfect the reporter plasmid into 293T cells. We then co-transfected the hsa-miR-329-5p mimics or nc into 293T cells. After 48 h, we used the Dual Luciferase Reporter Assay System (Promega, USA) to detect firefly and renilla luciferase activity by microplate reader (BioTek, USA). Finally, we used Read1:Read2 to represent fluorescence activity.

### Nude mouse model

Five-week-old female BALB/c nude mice were purchased from the BEIJING HFK BIOSCIENCE COMPANY and maintained under pathogen-free conditions. A total of 5 × 10^6^ THP-1 cells in 100 μl of Sodium Chloride Physiological Solution were injected into armpit of each mouse. After 12 days, the transplanted nude mice were randomly divided into two groups (n = 4 each). And then sicirc or siCtrl (Ribo, Shanghai, China) was directly injected into the implanted tumor at the dose of 5 nmol (in 10 ul Sodium Chloride Physiological Solutio) per mouse every 4 days. Tumor volume (V) was monitored by measuring the length (L) and width (W) with vernier caliper and calculated with the formula V = (L × W^2^) × 0.5. After each injection, the tumor size was measured. A total of four injections and five measurements were taken. The entire experimental protocol was conducted in accordance with the guidelines of the local institutional animal care and use committee.

### Statistical analysis

Statistical analyses were analyzed by GraphPad Prism 7.00 software (GraphPad software, La Jolla, CA, USA) and SPSS 25 software (IBM, USA). SPSS was used to analyze whether the data conformed to normal distribution and homogeneity of variance, and GraphPad Prism was used to analyze the data. PCR data were presented as mean ± standard error of the mean (SEM). Specific statistical methods are illustrated in the corresponding illustrations in the results. P value less than 0.05 was considered statistical significance.

## Results

### Expression of hsa_circ_0001947 and hsa_circ_0058058 in AML patients

To investigate the expression of circRNAs in AML patients, we used circRNAs microarray to determine the differential expression profile between six newly diagnosed (ND) AML patients without prior treatment and four healthy controls. The clustered heatmap or volcano plot filtering was used to indicate the differentially expressed circRNAs in the bone marrow of AML patients. A total of 147 upregulated circRNAs and 317 downregulated circRNAs were identified, which had aberrant expression more than two-fold in AML patients compared to healthy controls[21]. Based on the principle of log2 (fold changes) ≥1, P<0.05 and FDR<0.05, we selected two circRNAs, upregulated hsa_circ_0058058 and downregulated hsa_circ_0001947, as the candidate for the following study (Fig. 1A). The expression level of hsa_circ_0058058 or hsa_circ_0001947 in the individual AML patients or healthy control was shown by hierarchical clustering (Fig. 1B).

**Figure 1.**
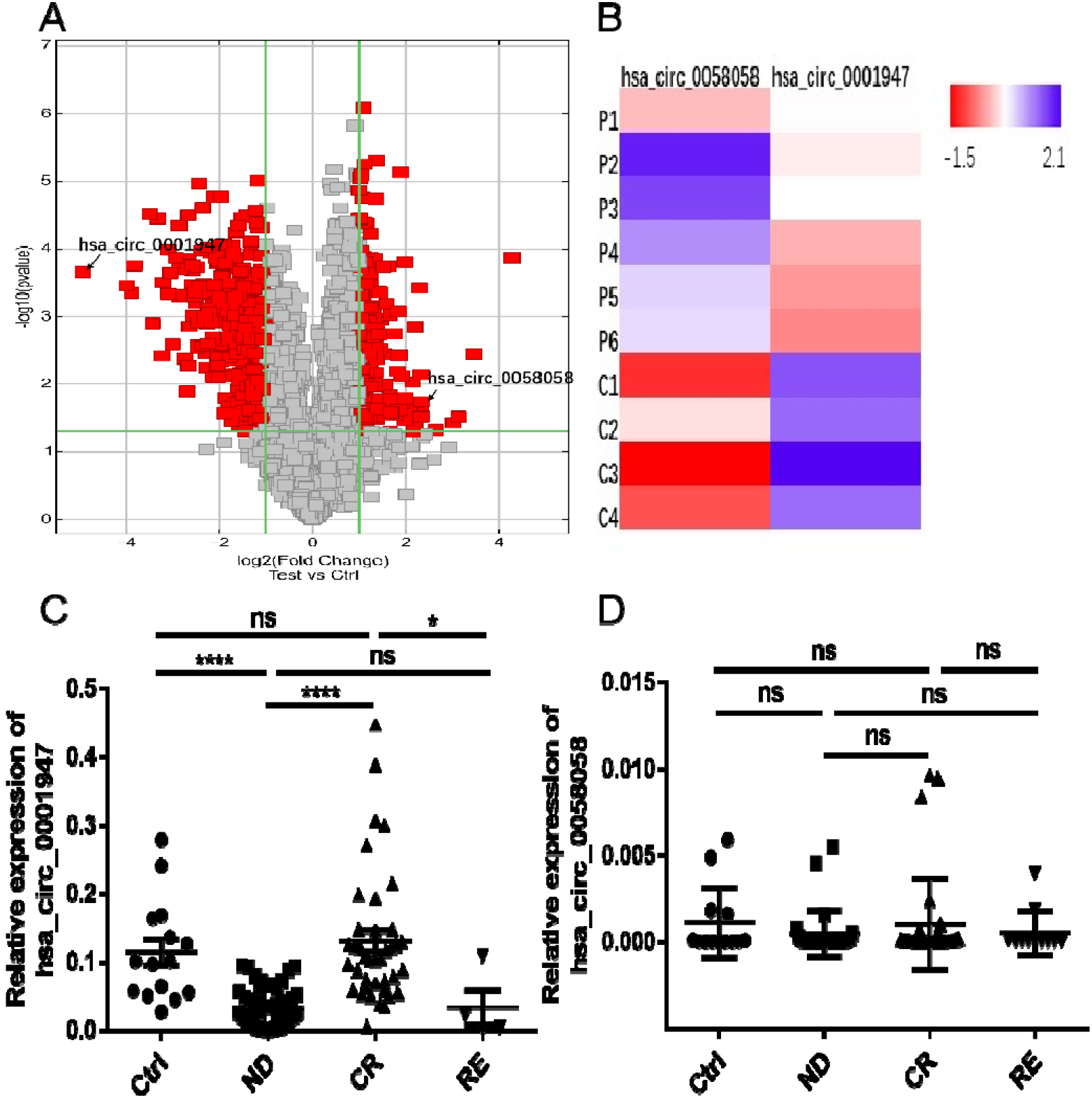
The microarray result of circRNAs in AML patients and controls and validation of hsa_circ_0058058 and hsa_circ_0001947 by qRT-PCR. (A). Volcano plot. The vertical lines correspond to two-fold changes up or down, respectively, and the horizontal line represents a p-value of 0.05. Red points in the plot represent the differentially expressed circRNAs with statistical significance. (B). Clustered heatmap for differentially expressed circRNAs. Rows represent indicidual patients or control and columns represent circRNAs. P, AML patients; C, controls. The color scale runs from red (low intensity) to white (medium intensity), to purple (strong intensity). (C). The expression of hsa_circ_0001947 in 102 AML patients (ND, n=59; CR, n=39; RE, n=4) and 15 controls. (D). The expression of hsa_circ_0058058 in 102 AML patients and 15 controls. * p < 0.05, ****p < 0.0001; ns, no significance. Data were analyzed using Mann Whitney test.

We further validated the microarray results of the differential expression of hsa_circ_0058058 and hsa_circ_0001947 in another cohort of 102 AML patients and 15 healthy controls by qRT-PCR. We found that the expression level of hsa_circ_0001947 in newly diagnosed (median 0.0245, 0.00073-0.0947) or relapsed-refractory AML patients (median 0.0133, 0.0018-0.1088) was much lower compared to that in healthy controls (median 0.1015, 0.0276-0.2793, p < 0.0001). Meanwhile, the expression level of hsa_circ_0001947 was elevated in patients achieved complete remission (median 0.1058, 0.0067-0.4475, p<0.0001) compared to that in the ND group (Fig. 1C). However, the expression of hsa_circ_0058058 showed no statistical difference between AML patients and controls (Fig. 1D). Therefore, hsa_circ_0001947 was selected for the following functional study.

### Characterization of hsa_circ_0001947 in AML

As the relationship between circRNA and linear RNA isoforms was complicated, we also compared hsa_circ_0001947 and its linear isoform. According to circBase database, we found that the spliced mature sequence of hsa_circ_0001947 was derived from the exon of *AFF2*, whose length is 861bp (Fig. 2A). Moreover, we confirmed the head-to-tail splicing structure in the qRT-PCR product of hsa_circ_0001947 with the expected size by Sanger sequencing (Fig. 2B). The stability of hsa_circ_0001947 and AFF2 linear mRNA was further tested, and our results showed that hsa_circ_0001947 harboring a loop structure was resistant to be digested by RNase R, while AFF2 mRNA could be degraded (Fig. 2C, D).

**Figure 2.**
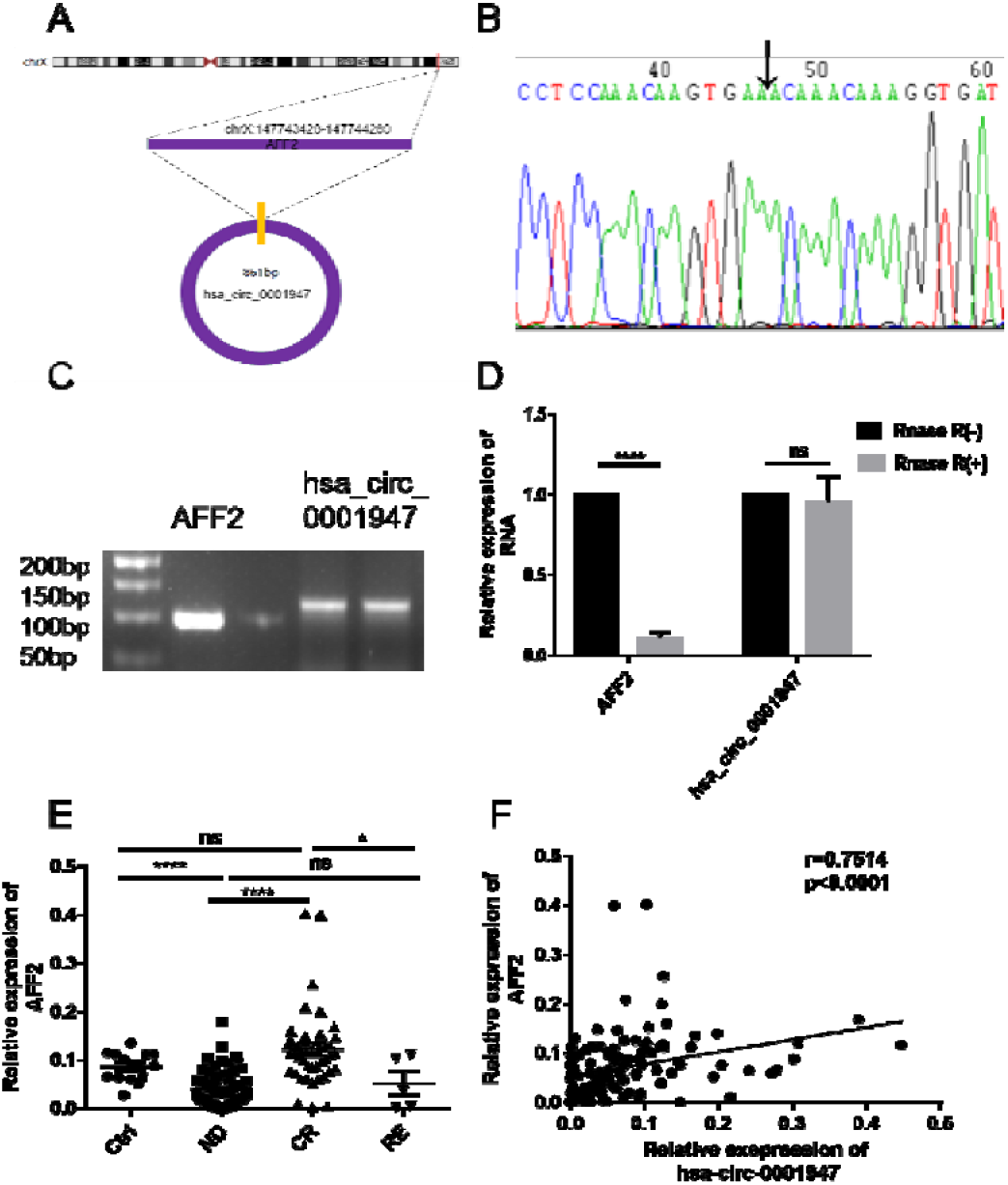
Identification of the circular structure and clinical features of hsa_circ_0001947. (A). The spliced mature sequence length of hsa_circ_0001947 derived from the AFF2 mRNA is 861 bp. (B) The results of Sanger sequencing from the head-to-tail splicing in the RT-PCR product of hsa_circ_0001947. (C, D). Expression levels of hsa_circ_0001947 and AFF2 mRNA in THP-1 cells treated with RNase R by qRT-PCR. Data were analyzed using Unpaired t test. (E). AFF2 mRNA expression in 15 healthy controls and 106 AML patients at different stages (ND, n=60; CR, n=41; RE, n=5). Data were analyzed using Mann Whitney test. (F). Correlation between the expression levels of hsa_circ_0001947 and AFF2 mRNA by qRT-PCR in healthy controls and AML patients at different stages. Data were analyzed using Pearson correlation. *p < 0.05, ****p < 0.0001; ns, no significance.

As little is known about AFF2 mRNA in AML, we also determined the mRNA expression of AFF2 in the same cohort of AML patients and healthy controls. Similar with hsa_circ_0001947, we found the mRNA expression level of AFF2 was lower in ND (median 0.0293, 0.0001-0.1276) or RE (median 0.0384, 0.0019-0.1118) AML group and higher in control (median 0.0915, 0.0281-0.1358) or CR (median 0.1134, 0.0017-0.4005) group (Fig. 2E). Furthermore, a positive correlation between hsa_circ_0001947 and AFF2 mRNA was observed in healthy controls or AML patients at different stages (Fig. 2F).

### Hsa_circ_0001947 is clinically favorable in AML patients

To explore the relationship between hsa_circ_0001947 and clinical practice, we analyzed the clinical significance of the hsa_circ_0001947 in AML patients. We found that the expression level of hsa_circ_0001947 was negatively correlated with white blood cell (WBC) (r=−0.369, P<0.05) and positively linked to hemoglobin (HGB) in AML patients (r=0.510, P<0.01) (Fig. 3A, B). Next, we analyzed the percentage of primitive cells in bone marrow (BM%) or peripheral blood (PB%). The results revealed that hsa_circ_0001947 expression level was negatively correlated with BM% or PB% in AML patients (r=−0.496, P<0.01; r=−0.429, P<0.05, respectively) (Fig.3C, D). Associations between hsa_circ_0001947 and several AML clinical features are shown in Table 3, demonstrating no significant difference in the age, gender, French-American-British (FAB) classification, which imply that hsa_circ_0001947 might function independently from those factors.

**Figure 3.**
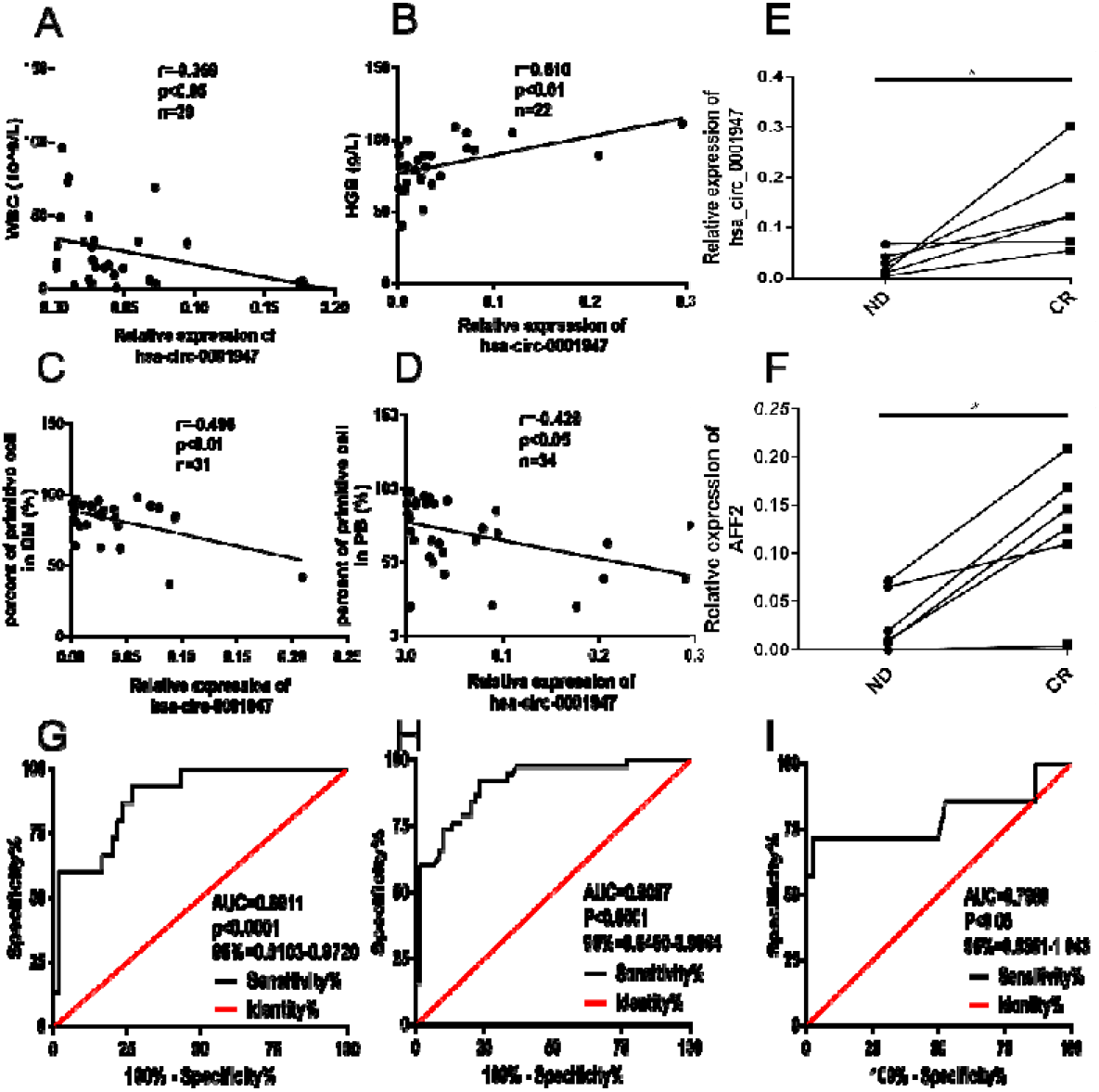
Clinical correlation of hsa_circ_0001947 in AML patients and ROC curve analysis of hsa_circ_0001947 and AFF2 mRNA in AML patients. (A, B, C, D) The results of correlation between the expression level of hsa_circ_0001947 and WBC, HGB, BM%, PB% in ND AML patients. Rows represent the relative expression of hsa_circ_0001947 and columns represent clinical features of hsa_circ_0001947 in AML patients. Data were analyzed using Pearson correlation. (E, F). Hsa_circ_0001947 and AFF2 mRNA expressions were measured in matched-pair samples acquired from six available follow-up AML patients at the time when they were at ND and CR stage. Data were analyzed using Wilcoxon test. (G H, I) Diagnostic values of hsa_circ_0001947 in different stages in AML patients. * *p* < 0.05.

To further investigate the influence of chemotherapy on hsa_circ_0001947, we followed the whole treatment process of six AML patients, who were treated with standard chemotherapy and then achieved CR. In matched-pair bone marrow samples acquired from those six AML patients at both ND and CR stage, a significant increase of hsa_circ_0001947 expression was demonstrated in CR stage compared with ND stage (Fig. 3E), indicating the restoration of dys-regulated hsa_circ_0001947 expression after chemotherapy. At the same time, a significant increase of AFF2 mRNA expression was also found in CR stage compared with their ND stage (Fig. 3F). These results suggested that hsa_circ_0001947 may be a biomarker for the diagnosis and prognosis in AML.

**Table 3.**
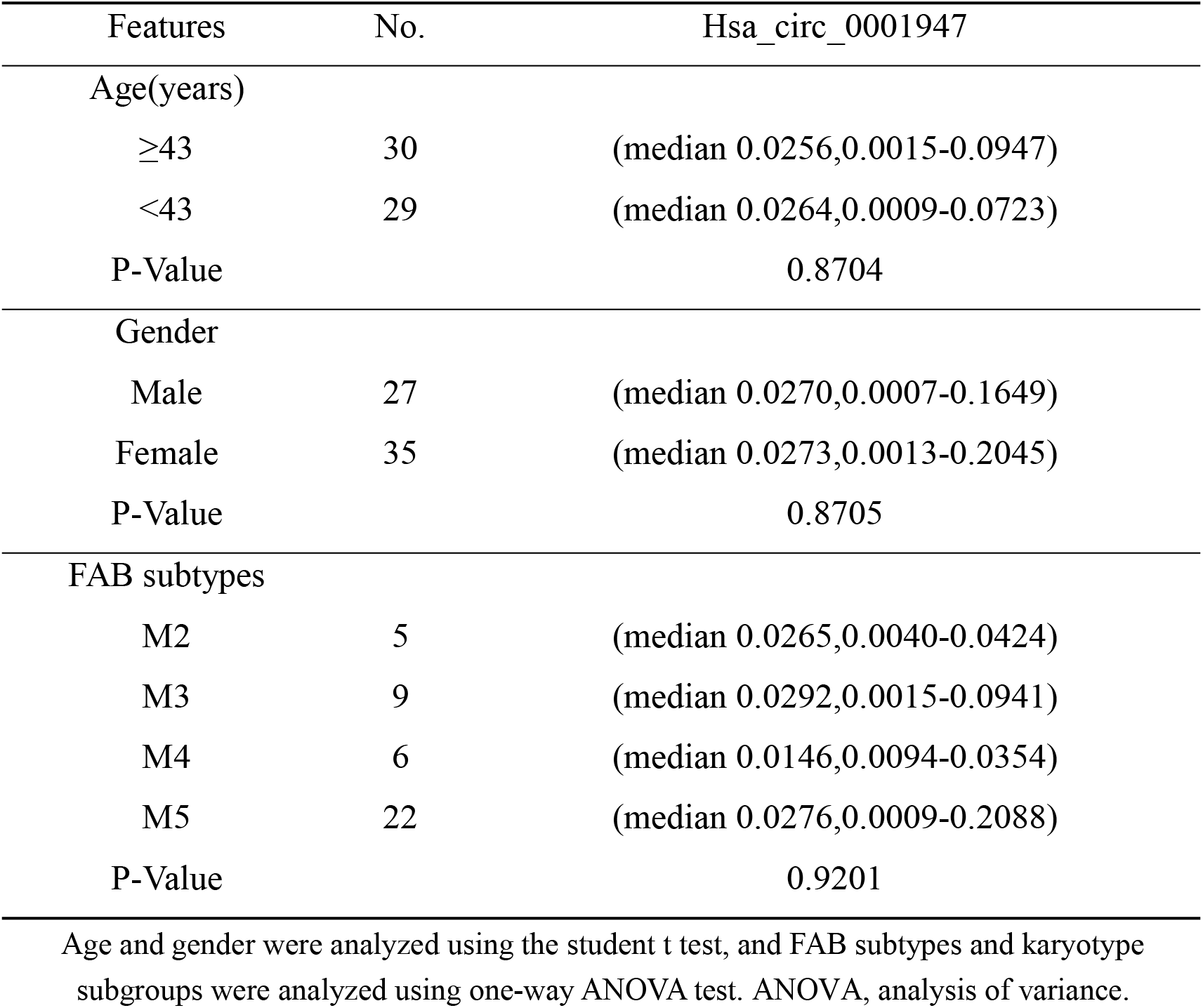
Expression of hsa_circ_0001947 in different subgroups of ND AML patients.

| Features | No. | Hsa_circ_0001947 |
| --- | --- | --- |
| Age(years) |  |  |
| ≥43 | 30 | (median 0.0256,0.0015-0.0947) |
| <43 | 29 | (median 0.0264,0.0009-0.0723) |
| P-Value |  | 0.8704 |
| Gender |  |  |
| Male | 27 | (median 0.0270,0.0007-0.1649) |
| Female | 35 | (median 0.0273,0.0013-0.2045) |
| P-Value |  | 0.8705 |
| FAB subtypes |  |  |
| M2 | 5 | (median 0.0265,0.0040-0.0424) |
| M3 | 9 | (median 0.0292,0.0015-0.0941) |
| M4 | 6 | (median 0.0146,0.0094-0.0354) |
| M5 | 22 | (median 0.0276,0.0009-0.2088) |
| P-Value |  | 0.9201 |

Age and gender were analyzed using the student t test, and FAB subtypes and karyotype subgroups were analyzed using one-way ANOVA test. ANOVA, analysis of variance.

### ROC curve analysis of hsa_circ_0001947 in AML patients

The area under the ROC curve (AUC) was used to determine the diagnostic value of hsa_circ_0001947 for AML. The AUC of hsa_circ_0001947 at the ND stage was 0.8911 (0.8103–0.9720, P <0.0001); the sensitivity and specificity were 93.33% and 73.33% (Fig. 3G). At the CR stage, the AUC of hsa_circ_0001947 was 0.9057 (0.8450–0.9664, P <0.0001); the sensitivity and specificity were 92.11% and 76.67% (Fig. 3H). Meanwhile, the AUC of hsa_circ_0001947 at the RE stage was 0.7989 (0.5551–1.043, P <0.05); the sensitivity and specificity were 71.43% and 97.37% (Fig. 3I). These results indicate that the expression difference of hsa_circ_0001947 make itself of significant diagnostic value.

### Downstream bioinformatics analysis of the Hsa_circ_0001947

Considering the results above, hsa_circ_0001947 is very likely to be a special AML-related circRNA. Based on the previous reports[22–24], we wondered whether hsa_circ_0001947 could also function as a miRNA sponge in AML and regulate its own circRNA-miRNA-mRNA network. With the help of TargetScan and Miranda software, we predicted top 3 of its most potential miRNA targets (in Fig. 4A). The results from Mirwalk3.0 (Based on TarPmiR prediction algorithm) showed that hsa-miR-329-5p, hsa-miR-10b-3p, hsa-miR-488-3p exhibited the most powerful circRNA-miRNA-mRNA gene interaction networks (Fig. 4B)[25]. Among hundreds of mRNA targets identified from sequence-pairing with those top three miRNAs, four of them were targeted by more than one miRNA: CREBRF, KLHL15, KRT12 and INTS9, which were chosen to be our main focus for further study.

**Figure 4.**
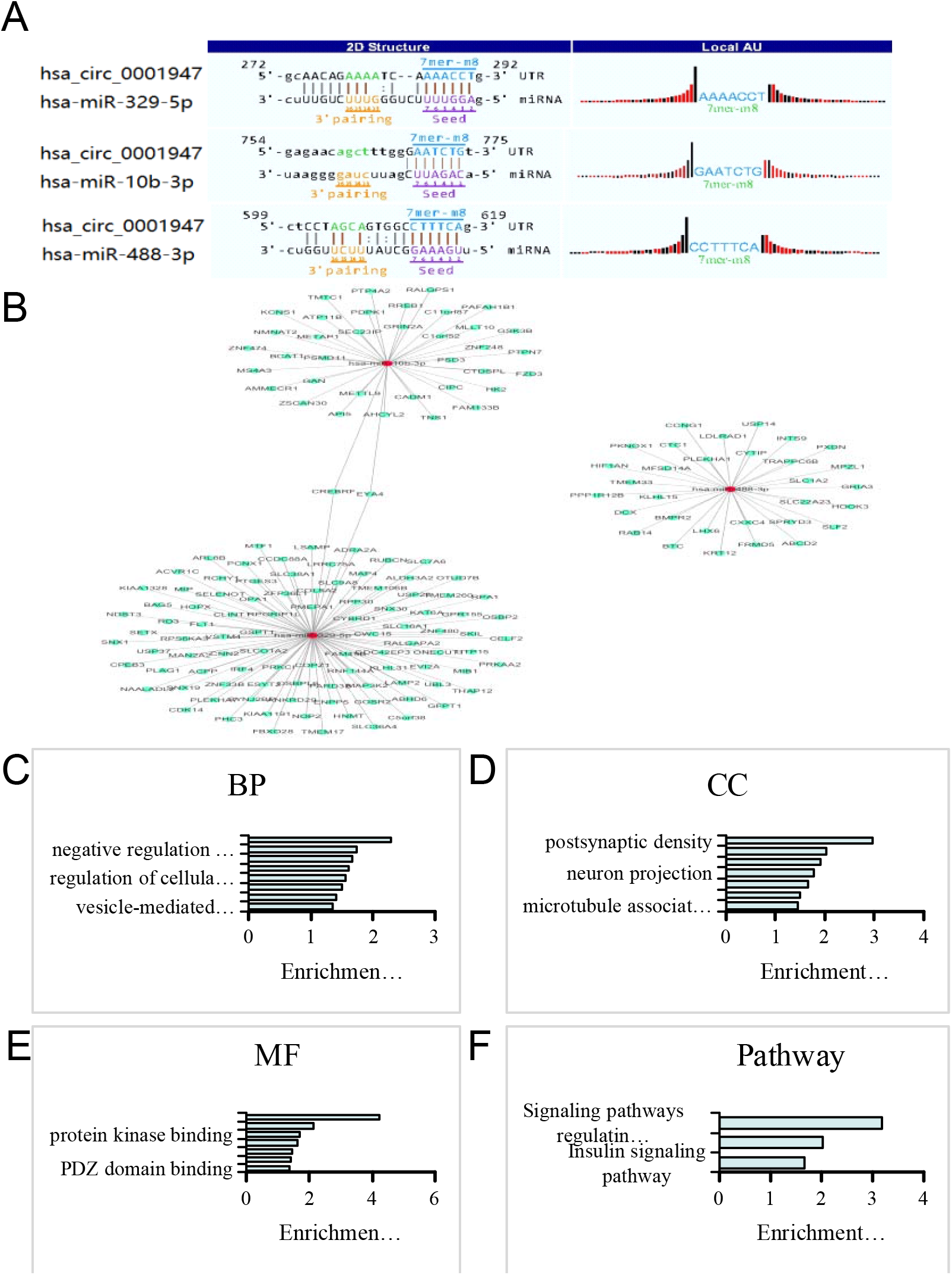
MiRNA prediction and downstream bioinformatics analysis of hsa_circ_0001947. (A) The 3 highest-ranking miRNAs matched hsa_circ_0001947. (B) CircRNA-miRNA-mRNA network of hsa_circ_0001947, focused on the three highest-ranking. (C, D, E) The results of the GO enrichment corresponds to the hsa_circ_0001947. Rows represent enrichment score and columns represent GO Term. (F). The results of the pathway related to hsa_circ_0001947. Rows represent enrichment score and columns represent pathway.

Besides, Gene Oncology (GO) and Kyoto Encyclopedia of Genes and Genomes (KEGG) pathway were used to investigate the functions of their target genes. Our results showed that the target genes related to hsa_circ_0001947 signature participated in various biological processes, such as multicellular organism growth and activation of protein kinase B activity (Fig. 4C). In addition, cellular component had also changing, such as postsynaptic density, cytoplasm and nucleoplasm (Fig. 4D). As shown in Fig. 4E, molecular function related to hsa_circ_0001947 signature is also exhibited, which consists of protein binding, phosphatidylinositol binding and protein kinase binding and so on. Furthermore, some vital pathways could also be influenced by the altered circRNAs, including insulin resistance, insulin signaling pathway, and signaling pathways regulating pluripotency of stem cells (Fig. 4F).

### Hsa_circ_0001947 manifests inhibitory effect for AML cells

To investigate the potential roles of hsa_circ_0001947 in AML cells, we conducted in vitro experiments by transfecting siRNAs (sicirc) or negative control (siCtrl) into THP-1 cells. The expression level of hsa_circ_0001947 was successfully downregulated after 48h transfection by qRT-PCR (Fig. 5A). Next, we detected the proliferation rate of THP-1 cell by Cell Counting Kit-8 (CCK-8) assay after transfecting with sicirc/siCtrl. The result showed that knocking down hsa_circ_0001947 could promote the proliferation of THP-1 cells compared with siCtrl (Fig. 5B). Then we dyed Annexin V and PI directly and found that the apoptotic frequency of cells after transfecting the sicirc was lower than that of the siCtrl after 48 h. (Fig. 5C). Further, the percentage of apoptotic cells were determined with Annexin V and PI after treatment with 2 uM cytosine arabinoside (Ara-c). Apoptotic cells after down-regulation of hsa_circ_0001947 was lower than that of control group (Fig. 5D). And then, in order to verify the candidate target gene, we conducted qRT-PCR in THP-1 cells transfecting with sicirc/siCtrl, and the results revealed that the downstream target gene CREBRF was also down-regulated after the hsa_circ_0001947 down-regulation, which was in accordance with our prediction (Fig. 5E).

**Figure 5.**
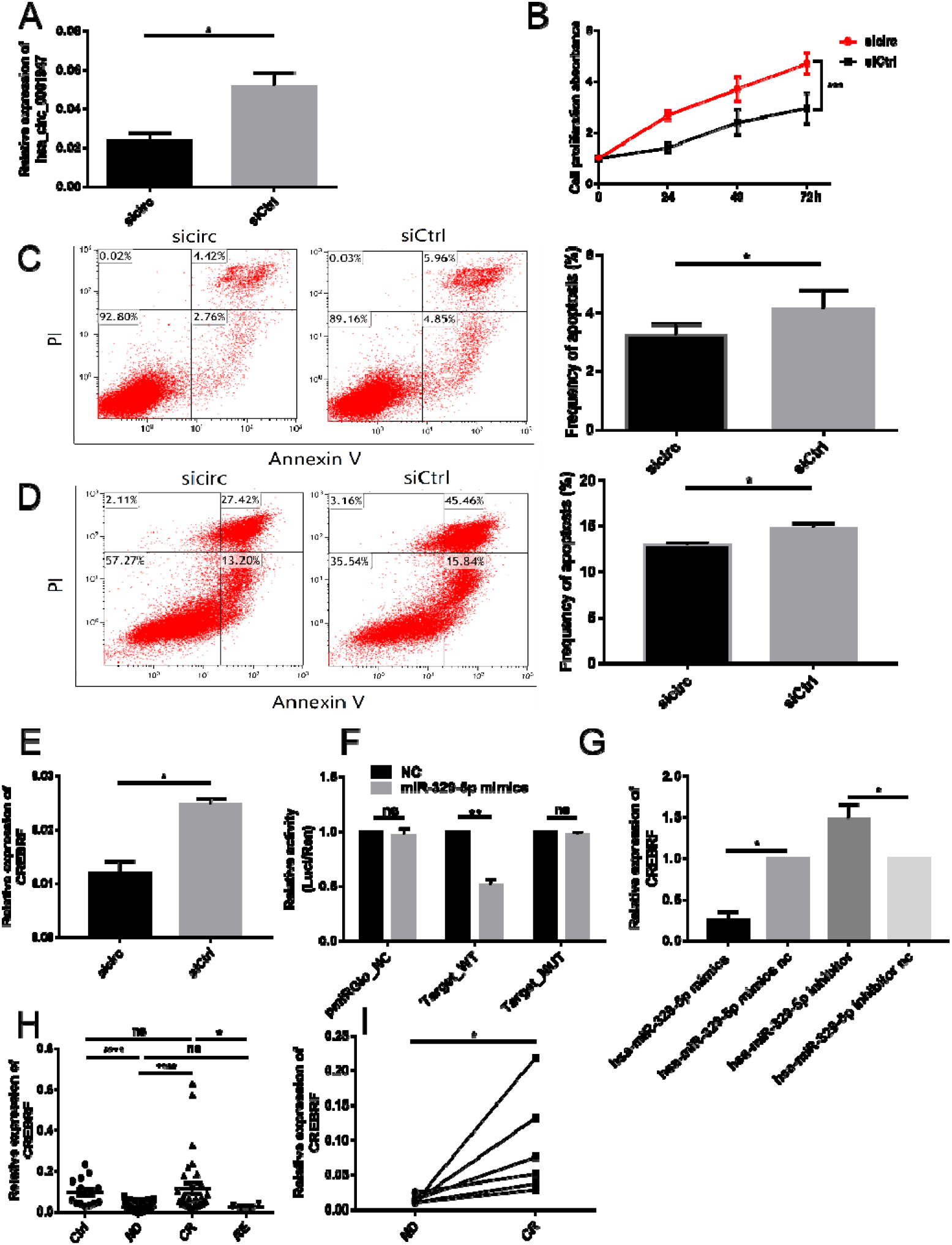
Hsa_circ_0001947 inhibits the proliferation, apoptosis in AML cell and validation of miRNA and target genes. (A). Verification of hsa_circ_0001947 downregulation efficiency by qRT-PCR. Data were analyzed using Mann Whitney test. (B). Proliferation of THP-1 cells were assessed by CCK-8 assays and proliferation rates at 24h, 48h, and 72 h were calculated. Data were analyzed using 2way ANOVA. (C) The results of apoptosis were measured by flow cytometry with dual staining of Annexin V and PI for the THP-1 cells transduced with sicirc or siCtrl. Data were analyzed using Unpaired t test. (D). THP-1 cells transduced with sicirc or siCtrl were cultured with Ara-c (2 uM, 48 h) and apoptosis was measured by flow cytometry with dual staining of Annexin V and PI. Data were analyzed using Unpaired t test. (E). The results of expression levels of candidate target genes CREBRF in sicirc/siCtrl THP-1 cells by qRT-PCR. Data were analyzed using Mann Whitney test. (F) The results of dual-luciferase reporter assay for hsa-miR-329-5p and CREBRF. Data were analyzed using Unpaired t test. (G) The results of expression levels of candidate target genes CREBRF by qRT-PCR after transfection with hsa-miR-329-5p mimics and inhibitor for 48 h. Data were analyzed using Mann Whitney test. (H) CREBRF mRNA expression in 14 healthy controls and 101 AML patients at different stages (ND, n=65; CR, n=32; RE, n=4). Data were analyzed using Mann Whitney test. (I) CREBRF mRNA expressions were measured in matched-pair samples acquired from six available follow-up AML patients at the time when they were at ND and CR stage. Data were analyzed using Wilcoxon test. * *p* < 0.05, ** *p* < 0.01, *** *p* < 0.001, ****p < 0.0001; ns, no significance.

Next, we performed a dual-luciferase reporter assay to determine the direct target gene of hsa-miR-329-5p based on their complementary sequences. A hsa-miR-329-5p fragment was constructed. Then, we co-transfected a hsa-miR-329-5p mimics with the reporter gene into 293T cells. A significant reduction in luciferase reporter activity was observed relative to co-transfection with control RNA (Fig. 5F).

What’s more, to further determine the direct target gene relationship between hsa-miR-329-5p and CREBRF, hsa-miR-329-5p mimics/nc and inhibitor/nc were transfected into THP-1 cells. After applying the mimics, CREBRF expression decreased compared with negative control, while after transfection with inhibitor, CREBRF expression increased compared with negative control (Fig. 5G). Furthermore, qRT-PCR was performed to explore the CREBRF expression level in AML specimens. Similar with hsa_circ_0001947 or AFF2, CREBRF mRNA expression level was lower in ND (median 0.0150, 0.0018-0.0708) or RE (median 0.0212, 0.0120-0.0412) AML patients compared with control (median 0.0811, 0.0263-0.2349) or CR (median 0.0515, 0.0120-0.6329) group (Fig. 5H). Moreover, in matched bone marrow samples acquired from six AML patients, a significant increase of hsa_circ_0001947 expression was demonstrated after achieving complete remission (Fig. 5I).

### Hsa_circ_0001947 inhibits AML cells proliferation and promotes apoptosis via hsa-miR-329-5p/CREBRF

To further understand the effect of hsa-miR-329-5p /CREBRF on AML cells, we transfected hsa-miR-329-5p mimics or inhibitor into Thp-1 cells and then we detected the proliferation process of THP-1 cell by Cell Counting Kit-8 (CCK-8) assay. The results showed that transfecting hsa-miR-329-5p inhibitor could suppress the proliferation of THP-1 cells compared with negative control. However, not much has changed when mimics was applied (Fig. 6A, B). Furthermore, the frequency of apoptotic cells after transfecting hsa-miR-329-5p mimics was found lower than that of the control group after treatment with 2 uM Ara-c for 48 h. Meanwhile, the percentage of apoptotic cells after transfecting hsa-miR-329-5p inhibitor was elevated than that of control group after treatment with 2 uM Ara-c for 48 h (Fig. 6C, D, E). These results suggest that hsa_circ_0001947 inhibits the proliferation and promotes apoptosis in AML cell. Based on our results above, a detailed circRNA-miRNA-mRNA (hsa_circ_0001947-hsa-miR-329-5p-CREBRF) interaction network was presented for hsa_circ_0001947, allowing us to better understand its underlying mechanisms for function in AML.

**Figure 6.**
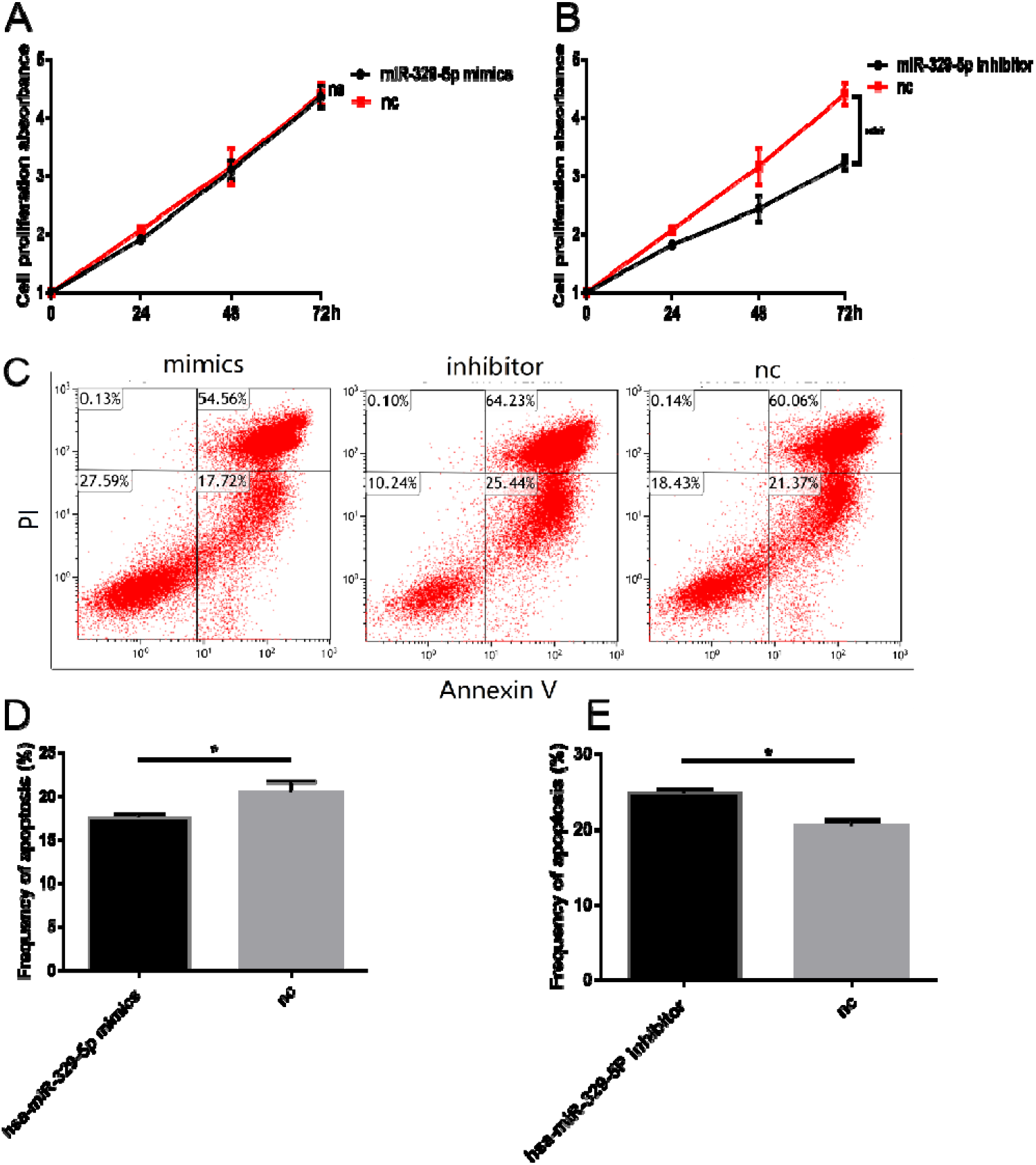
Hsa_circ_0001947 inhibits AML cells via hsa-miR-329-5p/CREBRF. (A, B). The hsa-miR-329-5p mimics or hsa-miR-329-5p inhibitor of proliferation in THP-1 cells were assessed by CCK-8 assays and proliferation rates at 24h, 48h and 72 h were calculated compared to negative control. Data were analyzed using 2way ANOVA. (C, D, E). THP-1 cells transduced with hsa-miR-329-5p mimics or hsa-miR-329-5p inhibitor were cultured with Ara-c (2 uM, 48 h) and apoptosis was measured by flow cytometry with dual staining of Annexin V and PI. Data were analyzed using Unpaired t test. * *p* < 0.05, *** *p* < 0.001; ns, no significance.

### Hsa_circ_0001947 inhibits AML cells growth in vivo

To explore the effects of hsa_circ_0001947 in vivo, THP-1 cells were injected into the nude mice unilateral axilla (n=4 mice in each group). Our results showed that the tumor volume in sicirc xenografts was bigger significantly than negative control xenografts (326.21 ± 285.95 mm^^3^ versus 190.96 ± 192.37 mm^^3^) and tumor weight were heavier in the sicirc group (Fig. 7A, B). Next, we checked the expression of hsa_circ_0001947 in tumor cells by RT-PCR. As shown in Figure 7C, the expression of hsa_circ_0001947 and the target gene CREBRF was successfully downregulated, as expected, but the linear gene AFF2 mRNA was not down-regulated in negative control. This also indicates that we specifically down-regulated hsa_circ_0001947. In addition, we also examined genes associated with proliferation and apoptosis. The expression of Bcl-2 and c-Myc were elevated in sicirc tumor than negative control. Moreover, Caspase-1 and Caspase-9 were reduced than negative control. Furthermore, the percentage of apoptotic cells from the sicirc xenografts was lower than the cells from negative control xenografts determined with Annexin V and PI (Fig. 7D). Next, we treated tumor cells with antibody Ki67 to compare the proliferation of the two groups. Results revealed that the proliferation ratio of the sicirc group was significantly higher than that of the negative control group (Fig. 7E).

**Figure 7.**
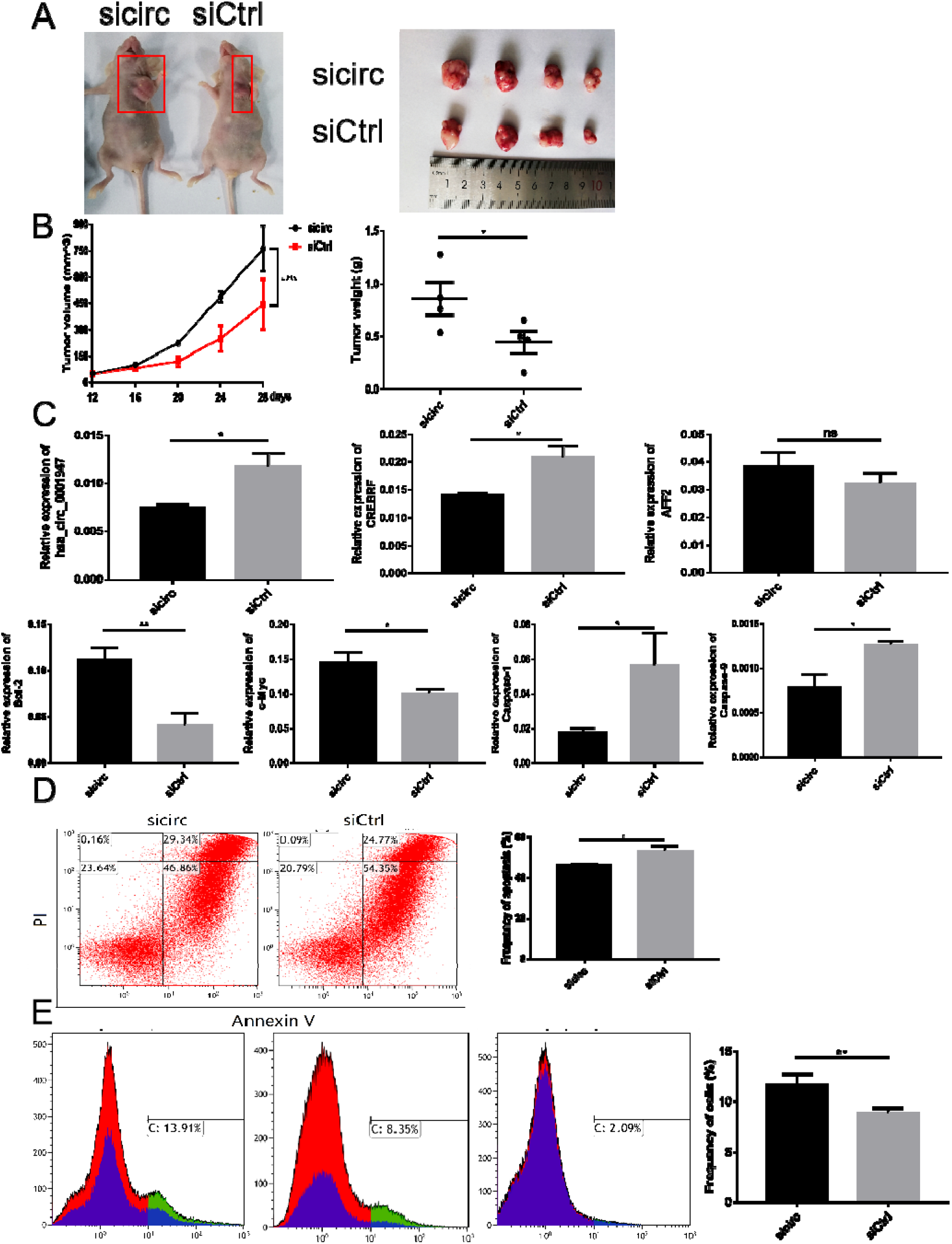
Downregulation of hsa_circ_0001947 in vivo. (A, B) The results of subcutaneous xenografts from the indicated THP-1 cell clones excised from nude mice tumor growth curve or tumor weight. Data were analyzed using 2way ANOVA or Unpaired t test. (C) The results of the expression of the related gene. Data were analyzed using Unpaired t test. (D) The results of the apoptosis for tumor cells from the sicirc xenografts and negative control xenografts measured by flow cytometry with dual staining of Annexin V and PI. Data were analyzed using Unpaired t test. (E) The results of THP-1 in sicirc xenografts and negative control xenografts measured by flow cytometry with staining of Ki67. Data were analyzed using Unpaired t test. * *p* < 0.05, ** *p* < 0.01; ns, no significance.

## Discussion

CircRNAs represent a widespread class of non-coding RNAs which are involved in the pathogenesis of many diseases[22, 26, 27], especially as important drivers of tumorigenesis or tumor suppressors in distinct human cancers[28]. CircRNAs which were regarded as miRNAs sponge have some characteristics in common. Through a splicing mechanism called backsplices, the precursor of the RNA is spliced, and the splicing donor connects to the upstream splicing receptor and forms circRNA[29]. These circRNAs are predominantly located in the cytoplasm, which occupies the same space of miRNAs[30]. CircRNAs with more predicting putative miRNAs binding sites are likely to play the role of ceRNA for miRNAs[27]. However, the function of circRNAs in the pathogenesis of AML has rarely been explored before. Therefore, we intend to explore the role of circRNAs for the identification of potential prognostic biomarkers or therapeutic targets for AML.

RNA sequencing or microarray analysis has been widely used to define the patterns of expressed circRNAs in various cancers. Lu et al. identified 1155 differentially expressed circRNAs from microarray analysis of four matched invasive ductal breast cancer and normal breast tissues[31]. Another study demonstrated that a total of 4266 circRNAs were differentially expressed in HCC and paired non-tumorous samples by the circRNA microarray analysis[32]. It was also reported that a panel of predicted circRNAs were identified by analyzing RNA sequencing data from 10 paired gastric cancer samples[18]. Our previous study has confirmed 147 up-regulated and 317 down-regulated circRNAs more than two-fold in AML patients compared to controls using high-throughout sequencing[21]. Subsequently, we focused on the differentially-expressed upregulated hsa_circ_0058058 and downregulated hsa_circ_0001947 based on the principle of log2 (fold changes) ≥1, P<0.05 and FDR<0.05. Then, according to the validation result, we identified hsa_circ_0001947, derived from the linear mRNA of *AFF2* exons (861 bp), as the candidate circRNA in AML. And we further demonstrated that AFF2 mRNA level was positively correlated with hsa_circ_0001947 in AML.

*AFF2*, extends over 600 kb in Xq27.3-q28, composed of 22 exons with a complex pattern of alternative splicing[33]. It has been demonstrated that *AFF2* mutations are associated with breast tumors[34]. Previous studies have reported that the AFF1-containing super elongation complex as a major regulator of development and cancer pathogenesis. In mammals, besides *AFF2*, the AFF family also includes two other members, AFF1 and AFF3. All of the family members share several conserved domains, including conserved N- and C-terminal domains, a serine-rich transactivation domain, and an ALF homology region[35]. GO enrichment analysis revealed that AFF2 mRNA was involved in various biological processes, such as cell adhesion, extracellular structure organization, collagen biosynthetic process and neuron development[36]. However, no studies revealed the role of *AFF2* and hsa_circ_0001947 in AML. In this study, we firstly confirmed the circular isoform of hsa_circ_0001947 by RNase R digestion test. Our result showed that hsa_circ_0001947 could still be detected by qRT-PCR under treatment with RNase R and demonstrated the stability of circRNAs, which is consistent with the previous reported study[6–8]. The expression of hsa_circ_0001947 has been demonstrated to be downregulated in AML patients and positively correlated with AFF2 mRNA expression, indicating that hsa_circ_0001947 might be associated with the development of AML.

CircRNAs are essential members of ceRNA networks, such as in the ‘circRNA-miRNA-mRNA’ axis, among which it functions as the ‘miRNA sponge’[7, 8]. This type of regulation is highly complicated because of the variability in the interactions between circRNAs and miRNAs. In our study, from the microarray analysis and downstream bioinformatics analysis of the hsa_circ_0001947, we chose hsa-miR-10b-3p, hsa-miR-329-5p and hsa-miR-488-3p as candidate miRNAs and CREBRF, KLHL15, KRT12 and INTS9 as candidate target genes. To further verify our prediction, we transfected mimics or inhibitor of predicted miRNAs into THP-1. The expression level of CREBRF, which is a target of hsa_circ_0001947, was found increased after inhibiting miRNAs. Luciferase assay further showed the precise and authentic interactions between circRNAs and miRNA, and miRNA and targeted genes. Finally, a detailed circRNA-miRNA-mRNA (hsa_circ_0001947-hsa-miR-329-5p-CREBRF) interaction network was presented for hsa_circ_0001947, allowing us to better understand its underlying mechanisms for function in AML. Collectively, these findings supported the hypothesis that hsa_circ_0001947 had an important role in AML progression.

What’s more, hsa-miR-329-5p gene has been predicted to be associated with apoptosis-related pathways (such as cell cycle and nucleotide excision repair pathway), metabolism-related pathways (such as biosynthesis of steroids, arginine and proline metabolism pathway), and immune-related pathways (such as toll-like receptor signaling pathway)[37]. For the first time we found that hsa-miR-329-5p not only had a high binding capacity with hsa_circ_0001947 but also demonstrated critical functions in AML progression. Then our findings suggested that hsa_circ_0001947 influenced the progression of AML by targeting CREBRF. Furthermore, some studies verified CREBRF participate in the regulation of angiogenesis by high-density lipoproteins[40]. In addition, it had been reported that CREBRF promotes the proliferation of human gastric cancer cells via the AKT signaling pathway[39]. And previous studies had reported that CREBRF was a potent tumor suppressor in glioblastoma by blocking hypoxia-induced autophagy via the CREB3/ATG5 pathway[38]. This is consistent with our research.

In summary, by combing circRNAs screening, functional verification together with clinical evidence, our study demonstrated the potential of hsa_circ_0001947 as promising prognostic biomarker in AML. What’s more, hsa_circ_0001947 functioned as a tumor suppressor by inhibiting cell proliferation in AML. Mechanistically, hsa_circ_0001947 acted as a hsa-miR-329-5p sponge restoring the expression of the hsa-miR-329-5p target gene CREBRF. This is the first report thoroughly investigating the biological functions of hsa_circ_0001947 and their clinical implications in AML, and potentially providing a new insight into the treatment of AML.

## Conclusions

In conclusion, our study reveals that hsa_circ_0001947 can serve as a novel potential biomarker for AML. Hsa_circ_0001947 inhibits the progression of AML by sponging hsa-miR-329-5p and further targeting CREBRF, thus providing a new insight into the treatment of AML.

## Abbreviations

AML: Acute myeloid leukemia
circRNAs: Circular RNAs
qRT-PCR: quantitative reverse transcription polymerase chain reaction
BMMCs: Bone marrow mononuclear cells
miRNA: microRNA
ceRNA: Competing endogenous RNA
ND: Newly diagnosed
CR: Complete remission
RE: Relapsed-refractory
WBC: white blood cell
HGB: hemoglobin
BM: Bone marrow
PB: peripheral blood
FAB: French-American-British
GO: Gene oncology
KEGG: Kyoto Encyclopedia of Genes and Genomes
CCK8: Cell Counting Kit-8
SEM: Standard error of the mean
ROC: Receiver-operating characteristic
AUC: Area under the ROC curve
GAPDH: Glyceraldehyde 3-phosphate dehydrogenase
Ara-c: cytosine arabinoside
ANOVA: analysis of variance.

## Acknowledgments

We sincerely thank all the participants in the study.

## Authors’ contributions

F.H. performed the experiments and analyzed data and wrote the manuscript. C.Z. collection of data. W.L. revised the paper. R.W. and C.Z. and X.Y. provided essential experimental tools. D.M designed the research, provided reagents and revised the paper. All authors read and approved the final manuscript.

## Availability of data and materials

All data generated or analyzed during this study are included in this published article and its supplementary information file.

## Ethics approval and consent to participate

In vivo studies were approved by the Institutional Animal Care and Use Committee.

## Consent for publication

Not applicable.

